# Phenotypic drought stress prediction of European beech (*Fagus sylvatica*) by genomic prediction and remote sensing

**DOI:** 10.1101/2023.03.29.534688

**Authors:** Markus Pfenninger, Liam Langan, Barbara Feldmeyer, Barbara Fussi, Janik Hoffmann, Renan Granado, Jessica Hetzer, Muhidin Šeho, Karl-Heinz Mellert, Thomas Hickler

**Affiliations:** Bayerisches Amt für Waldgenetik, Bayerisches Staatsministerium für Ernährung, Landwirtschaft und Forsten, Teisendorf, Germany; Department Molecular Ecology, Senckenberg Biodiversity & Climate Research Centre, Frankfurt/Main, Germany; Department Biogeography, Senckenberg Biodiversity & Climate Research Centre, Frankfurt/Main, Germany; Department Biogeography, J.W.Goethe University, Frankfurt/Main, Germany; Functional Environmental Genomics, LOEWE-Centre Translational Biodiversity Genomics, Frankfurt, Germany

## Abstract

Current climate change species response models usually not include evolution. We integrated remote sensing with population genomics to improve phenotypic response prediction to drought stress in the key forest tree European beech (*Fagus sylvatica* L.). We used whole-genome sequencing of pooled DNA from natural stands along an ecological gradient from humid-cold to warm-dry climate. We phenotyped stands for leaf area index (LAI) and moisture stress index (MSI) for the period 2016-2022. We predicted this data with matching meteorological data and a newly developed genomic population prediction score in a Generalised Linear Model. Model selection showed that addition of genomic prediction decisively increased the explanatory power. We then predicted the response of beech to future climate change under evolutionary adaptation scenarios. A moderate climate change scenario would allow persistence of adapted beech forests, but not worst-case scenarios. Our approach can thus guide mitigation measures, such as allowing natural selection or proactive evolutionary management.

## Introduction

While the anthropogenic greenhouse gas emission driving global warming continues more or less unabated ^1^, our understanding of climate change impacts on biodiversity and how to potentially mitigate its consequences, is not keeping pace ^2^. However, accurately predicting the outcomes of climate change for ecological keystone species, including evolutionary processes, is especially important because entire ecosystems and the benefits they provide to humanity rely on their persistence.

The European beech, *Fagus sylvatica* L., is playing a significant ecological role as beech forests provide a habitat for over 6,000 plant and animal species ^3, 4^. Beyond the key position in European woodland ecosystems, the importance of beech for ecosystem services, the economy and society can hardly be overestimated ^5^. Therefore, ancient and primeval beech forests of Europe are listed as UNESCO world heritage ^7^. This common dominant deciduous tree species of Central Europe out- competes other species due to its shade tolerance ^6^. Its distribution is mainly limited by water availability, as the tree cannot tolerate extended wet or dry conditions ^8^. Recent drought years have had a severe impact on the beech trees in Germany ^9^, severely damaging or killing locally up to 7% of trees. As previously shown ^10^, it was primarily medium to old-aged beeches that were affected by drought stress. However, this mortality might occur only many years after the actual drought event^11^. Accordingly, the German forest inventory data shows still increasing beech mortality, while those of most other assessed tree species has been declining again. However, local populations vary in tolerance to drought ^12, 13^, and the impact on individual tree vitality depends at least partially on their genomic composition ^14^. Because prolonged drought periods are predicted for the decades to come^15^, the species may severely suffer under future climatic conditions ^8^. Accurate predictions of of climate change impacts on beech forests are therefore urgently needed for the development of efficient mitigation strategies.

Several recent contributions identified evolutionary genomics as fast and efficient approach for accurate predictions of climate change impacts on species ^16–19^. These approaches suggest mainly genomic environmental associations (GEA) to identify genetic variation associated with environmental gradients and then predict either their spatial shifts or (mal)adaptation ^16^. These approaches rely on two crucial assumptions. First, it is supposed that a linear or continuous relation between the frequency of the genetic variants underlying the trait and the selectively relevant environmental variation exists ^17^. However, it has recently been shown that the variants underlying a continuous, additive, quantitative polygenic trait need not follow a clinal pattern ^18^. This is the case even if the trait perfectly co-varies with the selectively relevant environmental gradient, because intermediate phenotypes of even moderately polygenic traits can arise from thousands of different genotypes. Consequently, the expectation of continuous allele frequencies along environmental gradients may lead to the identification of false positives^18^. As most traits have a polygenic basis ^19^, this finding appears to be a serious challenge for GEA.

The second critical assumption is that the populations screened are locally adapted to their position on the environmental gradient ^16^. This assumption may be violated, because anthropogenic climate change is ongoing, and depending on evolutionary potential, life-span etc., the populations currently assessed may be already mal-adapted ^20^. This could be especially true for long-lived species actually experiencing climate change instead of weather dynamics. A lack of local adaptation may arise from the partial management many species of human interest, like forest trees or otherwise exploited species, are subjected. Local populations could thus have been restocked, planted, selectively harvested or otherwise managed not allowing them to adapt to local conditions ^21^. This could be particularly relevant for beeches, as many current forests have emerged from afforestation ^22^.

The prediction of climate change responses of phenotypic key traits may be an alternative approach. The variation in most quantitative traits has both an environmental and a genetic component ^23^. If the genetic basis of the trait is known, a validated polygenic or genomic prediction score composed of the weighted sum of associated alleles can be calculated from the respective multilocus genotype at the identified trait loci ^24^. If estimates of phenotypic reaction norm and a genomic prediction score are available, prediction of the trait response under not yet observed environmental conditions should be possible ^25^.

We, propose here a climate change prediction approach for *F. sylvatica* based on genomic prediction (GP) of drought resistance as a key phenotypic trait drawing on the recent identification of the genomic basis of drought resistance in beech ^14^. We extend the genotype-based GP model to population allele frequency data. We include relevant weather observations for the period 2016- 2022 as environmental data to validate the model and explain observed phenotypic data from genotyped beech stands obtained by remote sensing for 15 beech populations across Germany. This validated model was then used to predict drought response to future climate scenarios under three different evolutionary adaptation scenarios, i) no adaptation at all, ii) observed rate of adaptation among growth classes and iii) evolution of maximum observed drought resistance (Figure 1).

**Figure 1.**
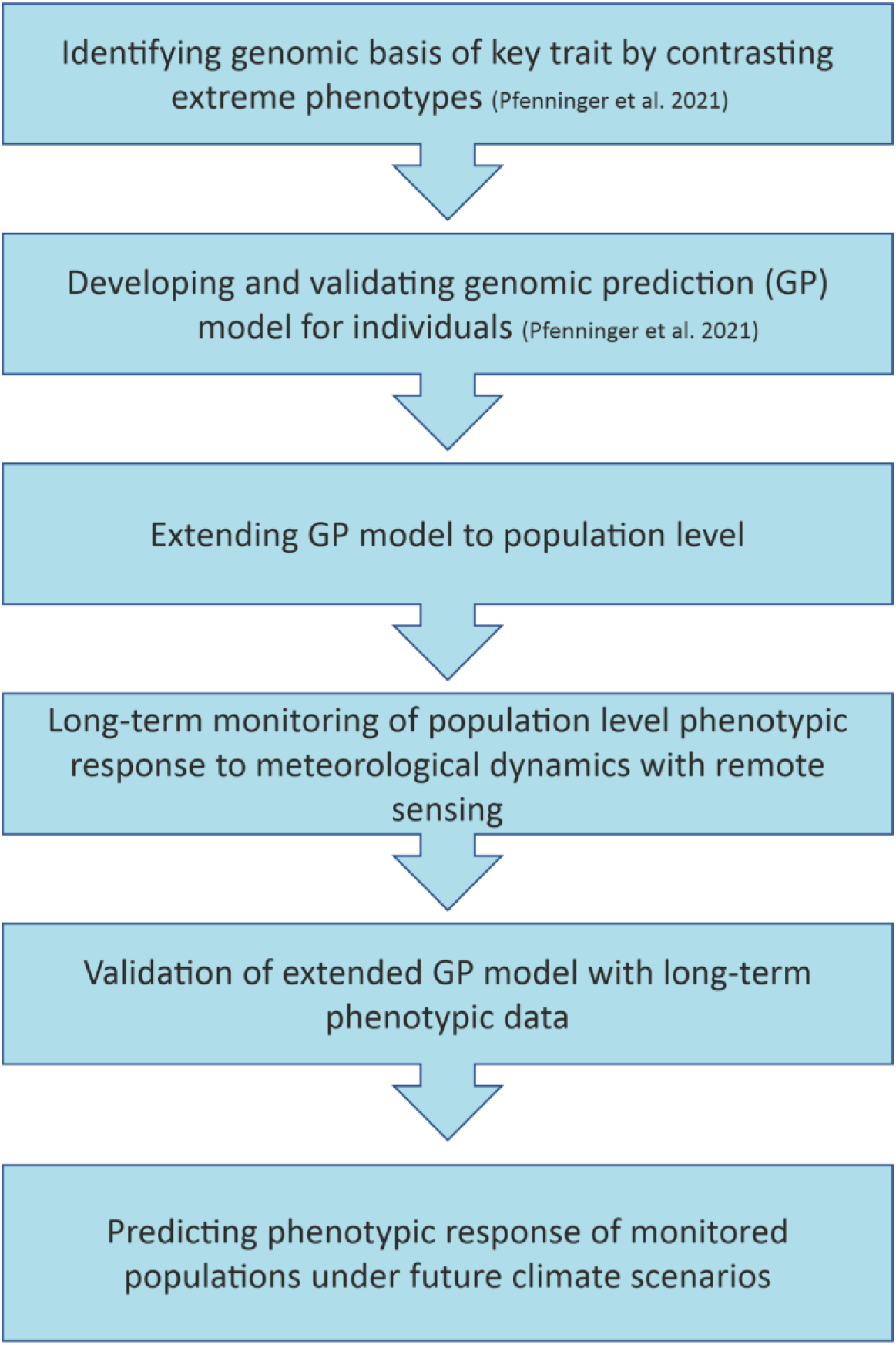
Conceptual proceeding.

## Results

### Long-term climate and weather

PCA on long-term climate data (1970 – 2000 ^27^) during the vegetation period (Apr - Sep) ordered the sampling sites (Table 1) along two meaningful axes. PCA1 (69.7% of variance) opposed warmer and cooler sites, while PCA2 (17.6%) distinguished between wet and dry sites (Figure 2b). Since the difference in precipitation was not substantial among the warmer sites, we applied PCA1 as predictor for summer drought risk and thus local adaptation.

**Table 1.** Sampling sites.

| site | location name | federal state | origin | Latitude | Longitude |
| --- | --- | --- | --- | --- | --- |
| CUN | Cunnersdorf | Sachsen | autochthon | 50.8462 | 14.1366 |
| KAR | Karlstadt (Arnstein) | Bayern | autochthon | 49.9659 | 9.7541 |
| KAU | Kaufbeuren | Bayern | unknown | 47.9155 | 10.5812 |
| KST | Königstein (Romberg) | Hessen | autochthon | 50.1917 | 8.4575 |
| LAN | Langen | Hessen | planted | 49.9964 | 8.6398 |
| LBA | Langenbrettach | Baden-Württemberg | unknown | 49.1991 | 9.3584 |
| MAR | Marxzell | Baden-Württemberg | autochthon | 48.8354 | 8.4239 |
| MOG | Mockrehna (GB) | Sachsen | unknown | 51.5614 | 12.7466 |
| MOM | Mockrehna (marginal) | Sachsen | unknown | 51.5625 | 12.7485 |
| OES | Östringen | Baden-Württemberg | autochthon | 49.1979 | 8.7473 |
| OTT | Ottendorf | Sachsen | autochthon | 51.2196 | 13.8383 |
| SEI | Seilershausen | Bayern | autochthon | 50.0721 | 10.4299 |
| TAU | Tauberbischofsheim | Baden-Württemberg | unknown | 49.6064 | 9.6542 |
| ULM | Ulm | Bayern | planted | 48.4559 | 10.066 |
| WUR | Wurzelberg | Thüringen | planted | 50.5029 | 11.035 |

**Figure 2.**
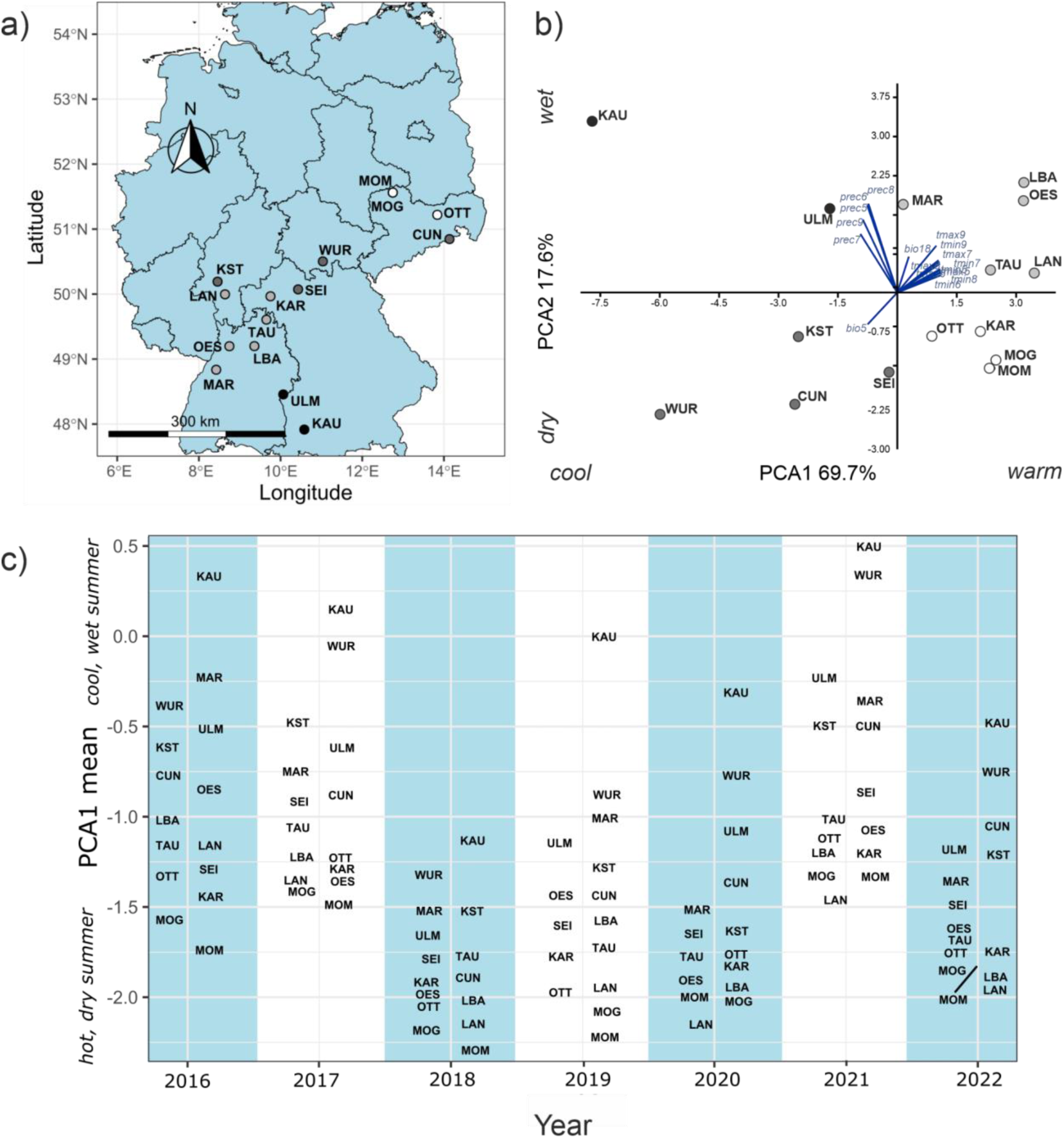
A) Geographic distribution of sampling sites, B) PCA of long-term climatic data during the vegetation period for the sampling sites, C) Annual means of PCA-scores on meteorological data during the vegetation period (May-Sep) for the sampling sites for each year 2016-2022. Higher values indicate cooler and moister years.

Weather data from DWD varied strongly during the observation period (2016-2022) from year to year and from site to site, as the plot of the annual PCA1 site score means (89% of variation) showed (Figure 2c). The summers of 2016, 2017 and 2021 were cool and wet and therefore served as baseline in analyses. The vegetation period in the years 2018, 2019, 2020 and 2022 were characterised by drought, i.e. high temperatures and little precipitation.

### Genetic variation and population structure

We obtained a mean sequencing coverage of 40X from 15 population pools with 48 canopy forming individuals each, i.e. 720 individuals. We scored about 2.6 million high-quality SNPs. Mean nucleotide diversity π within sites was 0.0080 (s.d. 0.0007) corresponding to one SNP per 125 base pairs. Mean theta ϴ was 0.0086 (s.d. 0.0008). Assuming a seed-to-seed mutation rate in the order of 2×10^-9^ gave an estimated effective population size N_e_ of about 10^6^. Mean F_ST_ of all pairwise comparisons in 33,768 1kb windows was 0.039 (s.d. 0.009). This was in the range of different age classes within two of the sites (LAN and KST, Figure 3a). Correlating the genome-wide mean F_ST_ among sampling sites with the geographic distance yielded no indication for isolation-by-distance (Mantel’s test r = -0.11, p = 0.786, Figure 3b).

**Figure 3.**
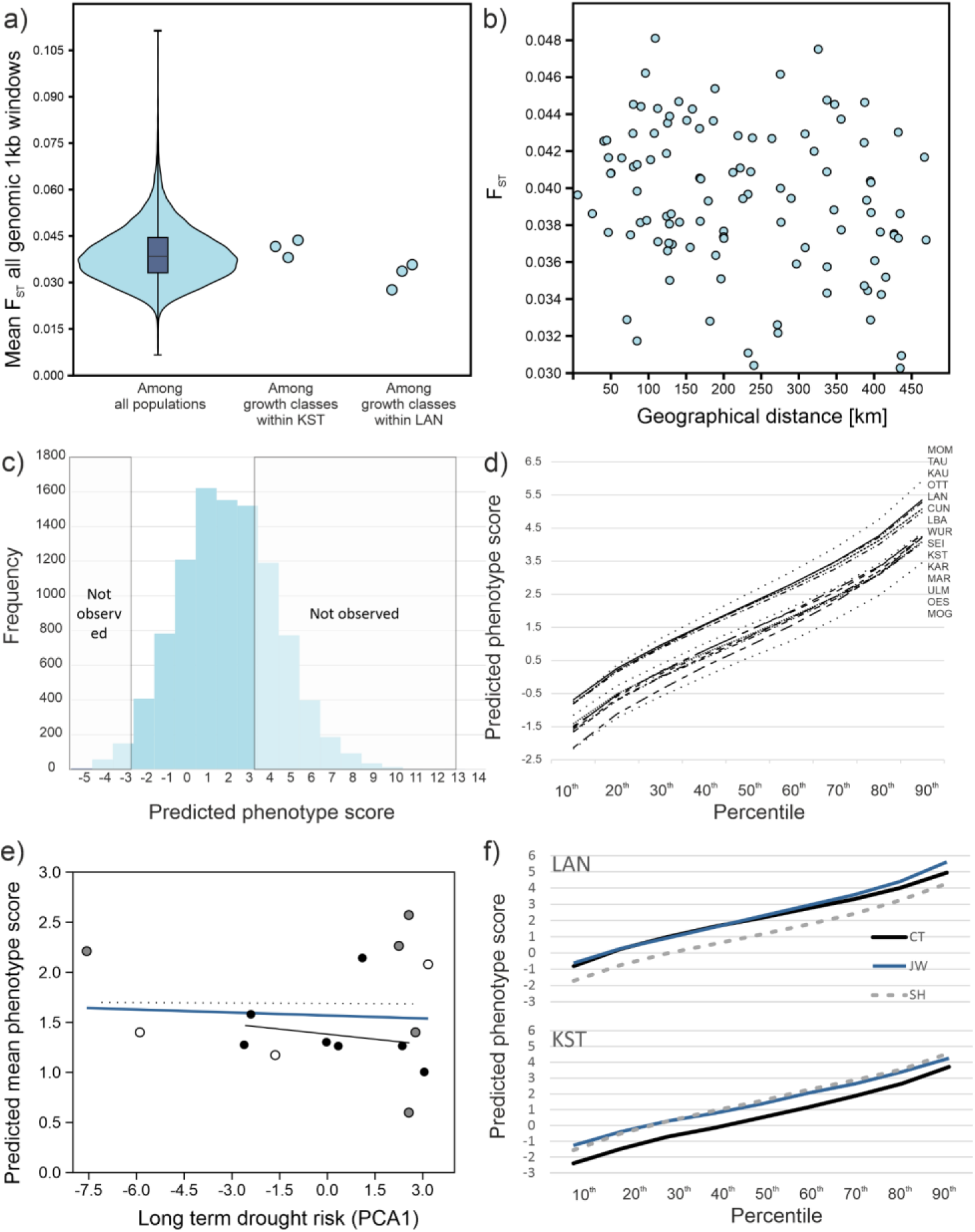
Population genomics and genomic prediction results. A) Violin plot of distribution of all site pairwise FSTs, compared to the F_ST_s among growth classes at the sites KST and LAN. B) Site pairwise FSTs against geographical distance between them. Mantel’s test showed no significant relation between FSTs and geographical distance (r = -0.226, p = 0.34). C) Frequency distribution of predicted phenotype scores of 10,000 simulated individuals from overall allele frequencies at 46 predictive loci. Individuals with predicted phenotype scores below -2.83 and above 3.3 (within dashed rectangles) were not observed in nature. D) Cumulated percentile distributions for genomically predicted phenotype distributions of the canopy forming beeches at the sampling site. Lower values indicate a larger proportion of drought resistant phenotypes. E) Plot of long term drought risk against PDPS. The blue line represents the overall linear regression, the black line the relation for autochthonous stands (black points) only, the dashed line for planted (white points) and unknown (grey points). F) Comparison of predicted phenotype distributions among growth classes at the sites LAN (above) and KST (below). The black lines indicate canopy forming trees, the grey dashed lines established trees not yet reaching the canopy and the blue line undergrowth below 1 m height.

### Genomically predicted population phenotypes

We used estimated allele frequencies with the genotypic prediction weight matrix of 46 trait associated SNP loci ^14^ to predict the phenotypes of 10,000 individuals from their simulated genotypes. The predicted drought phenotype score (termed PDPS hereafter) ranged from 0.599 (MOG) to 2.572 (MOM) with an average of 1.569 (s.d. 0.555, Figure 3d, Supp. Fig. 1). Distributions for all assayed sites were more or less normally distributed, which made the mean a useful statistic. Assuming that individuals with a phenotypic score of zero or less are resistant to drought conditions, the expected proportion of non-susceptible trees ranged from 25 – 45%, with a mean of 32.8%. The PDPS distribution based on the mean allele frequencies over all sites had a mean of 1.556 (s.d. 2.359) and ranged from -7.02 to 13.43. The realised predicted scores of 200 phenotyped and genotyped individuals ^14^ showed that scores below -2.83 and above 3.3 were not observed in the natural sample (Figure 3c).

Three of the 46 predictive loci showed a significant correlation (r > 0.5) of allele frequencies with either PDPS or drought risk score (Supp. Fig. 2). However, 445 out of 29,774 (1.5%) randomly chosen, highly differentiated (minimum allele frequency difference of 0.6 between two or more sites), unlinked SNPs showed a correlation of r >= 0.5 with the environmental gradient and were thus potential false positives.

### Local adaptation to drought risk

There was no correlation between PDPS and the long-term summer drought risk at the sites (r = 0.085, p = 0.93, Figure 3e). PDPS at the autochthonous stands alone did also not conform to expectations of local adaptation (Figure 3e). The patterns of PDPS distributions among growth classes differed. At the LAN site, the youngest (JW) growth class and the oldest class, the canopy forming trees (CT) showed very similar distributions except for the most drought-susceptible trees. The intermediate (SH) class had much more drought resistant predicted phenotypes. Its PDPS was one unit or 0.41 Haldanes (a shift in population mean expressed in standard deviations) smaller (Figure 3f). At the KST site, the CT class was the most drought resistant, while the younger classes showed more susceptible phenotype distributions. Over all classes, PDPS level at the KST site suggested a higher predicted drought resistance (Figure 3f).

### Results from remote sensing

Remote sensing yielded 955 measurements of canopy leaf area index (LAI) and moisture stress index (MSI) in the vegetation periods 2016-2022 for the 15 sites (∼9 observations per year and site). The overall observed mean LAI was 2.66 (s.d. 0.50) with a range from 1.28 to 4.80 and 0.67 (s.d. 0.07, range 0.29 – 1.02) for MSI. The means per vegetation period differed systematically among years and among sites for both parameters (Figure 4a+b). Within each year, mean LAI tended to decline over the vegetation period, independent of weather conditions (mean decrease between maximum and minimum = 0.98, s.d. = 0.51, Figure 4c). An increasing, but weaker trend was observed in MSI means (Figure 4d).

**Figure 4.**
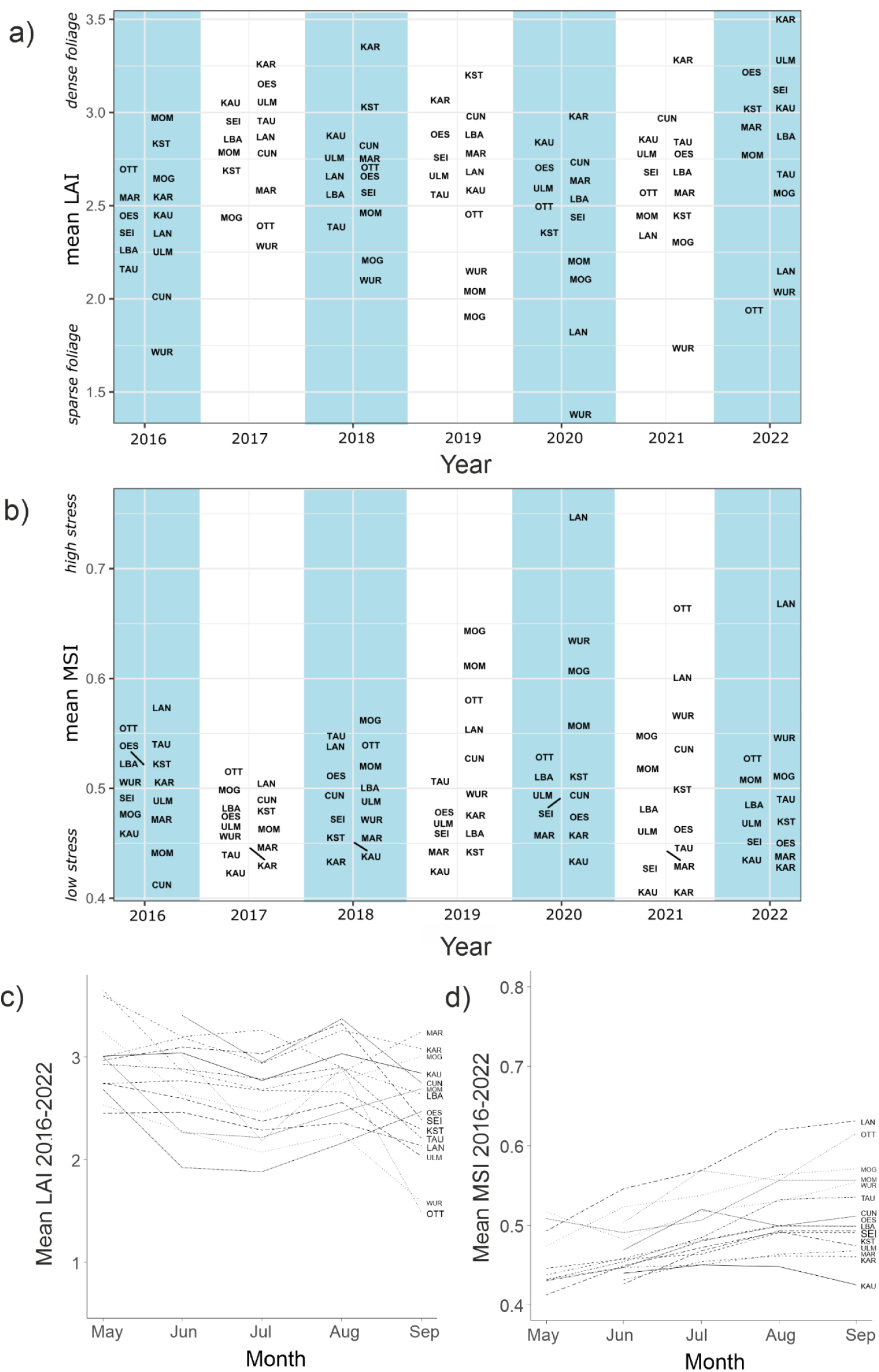
A) Stand-wide mean Leaf Area Index (LAI) data from Sentinel2 satellites during the vegetation period for the sampling sites in the observation period 2016-2022. Higher values indicate denser foliage. B) likewise for Moisture Stress Index (MSI). Higher values indicate more water stress. C) Mean temporal trajectory of LAI from May to September during the period 2016-2022. D) Likewise for MSI.

During 2016 and 2017, serving as non-drought baseline, the 1% quantile LAI for all individual pixels was 1.4, suggesting that values below this threshold were likely drought induced. Over the observation period, almost a third of pixels (0.328, s.d. 0.321) at all sites showed LAI < 1.4 at least once. The sites were highly differentially impacted (range 0.02 (KST) to 0.93 (WUR)). Pixels which showed an LAI of less than 1.4 for two or more years in a row, an MSI of 1.2 or more was measured in the first year, suggesting that this threshold induced long-term damage.

### Inference of fitness-relevant drought stress response thresholds

Satellite images and spatial projections of LAI and MSI revealed that the response of beech trees to the same drought conditions differed strongly within site (Figure 5 a-c). While the majority of pixels was not drought affected, retaining high LAI and low MSI values, few strongly affected trees over-proportionally influenced respective stand means.

**Figure 5.**
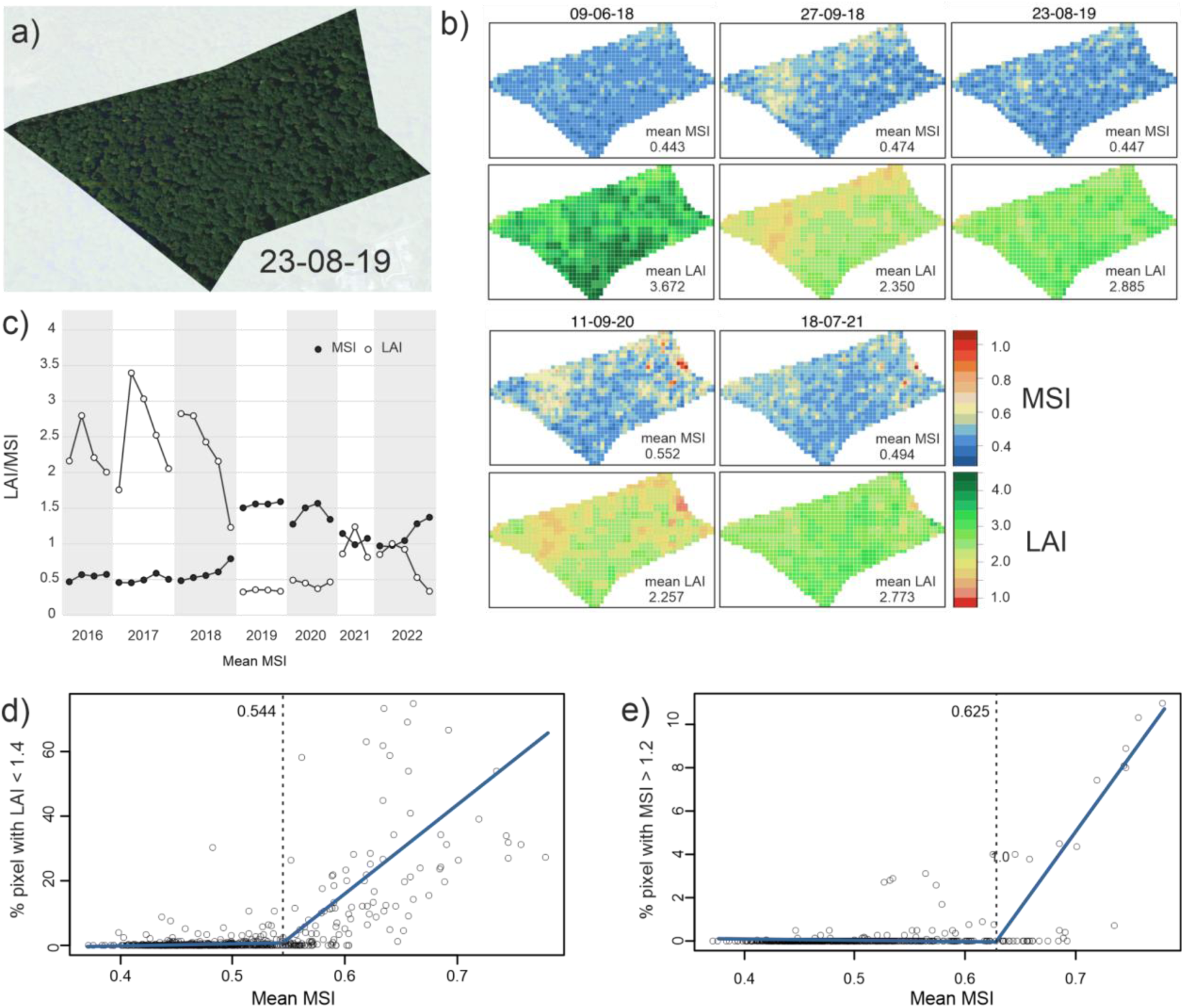
Damage threshold inference. A) High resolution satellite image of an exemplary site (KST) B) Spatial visualisation of MSI (above) and LAI (below) at KST in pixels of 10 x 10 m for a number of chosen dates. C) Exemplary temporal trajectory of LAI (white) and MSI (black) of a single pixel at site LAN. LAI and MSI became correlated only after the 2018 drought. E) Plot of stand-wide mean MSIs against the proportion of pixels in all stands with a LAI below 1.4. Values below this threshold indicate drought-induced leaf-loss. The blue line indicates the best fit of a segmented regression, the dashed line shows the inferred break-point (0.544). F) Plot of stand-wide mean MSIs against the proportion of pixels in all stands with a MSI above 1.2. Above this value, long-term leaf-loss occurs in the respective pixels. The blue line indicates the best fit of a segmented regression, the dashed line shows the inferred break-point (0.625).

The temporal trajectory of an exemplary single pixel showed that MSI remained about constant in 2016 and 2017 (non-drought baseline), as well as in early 2018 (Figure 5d). During the same period, LAI varied independently of MSI according to the seasonal pattern (Figure 5d). However, as MSI rose over a certain threshold during the 2018 drought, MSI and LAI started to covary (Figure 5d).

Plotting the percentage of pixels with LAI < 1.4 against mean MSI revealed a significant change in slope (Figure 5e). Segmented regression provided a significantly better fit to the data than a linear model (t = 30.496, d.f. = 876, p < 2.2 x 10^-16^). At 0.544 (s.e. 0.005), the slope estimate increased from 5.1 (s.e. 6.9) to 214 (s.e. 10.1). A mean MSI threshold of 0.625 marked the onset of longer lasting drought damages (%pixels with MSI > 1.2). Below this threshold, the slope was -0.44 (s.e. 0.41), above 51.8 (s.e. 2.22, slope change t = - 5.464, d.f. = 718 p < 6.4 x 10^-8^).

### Model selection of phenotypic drought response model

A GLM with the explanatory variables soil moisture, mean daily temperature in the month of observation, mean daily maximum temperature and the day of the year provided the best model fit for both response variables (LAI, MSI) with an ΔAIC of more than 10 units compared to the next best model (Table 2). Adding PDPS as variable explained the variance in LAI substantially better (ΔAIC = - 3.63, Table 2). Environmental variables cumulatively explained 67.1% of variance, genomic prediction 11.4%, while 21.5% remained unexplained. For MSI, inclusion of PDPS made the model decisively better (ΔAIC = -11.2, Table 2). Genomic prediction explained 14.8%, while environmental variables accounted for 65.9%, leaving 19.3% unexplained. Of this residual variation, 29.5% (= 5.7% of total variance) was explained by fixed differences among sampling sites (ANOVA: F =14.89, p = < 0.001).

**Table 2.**
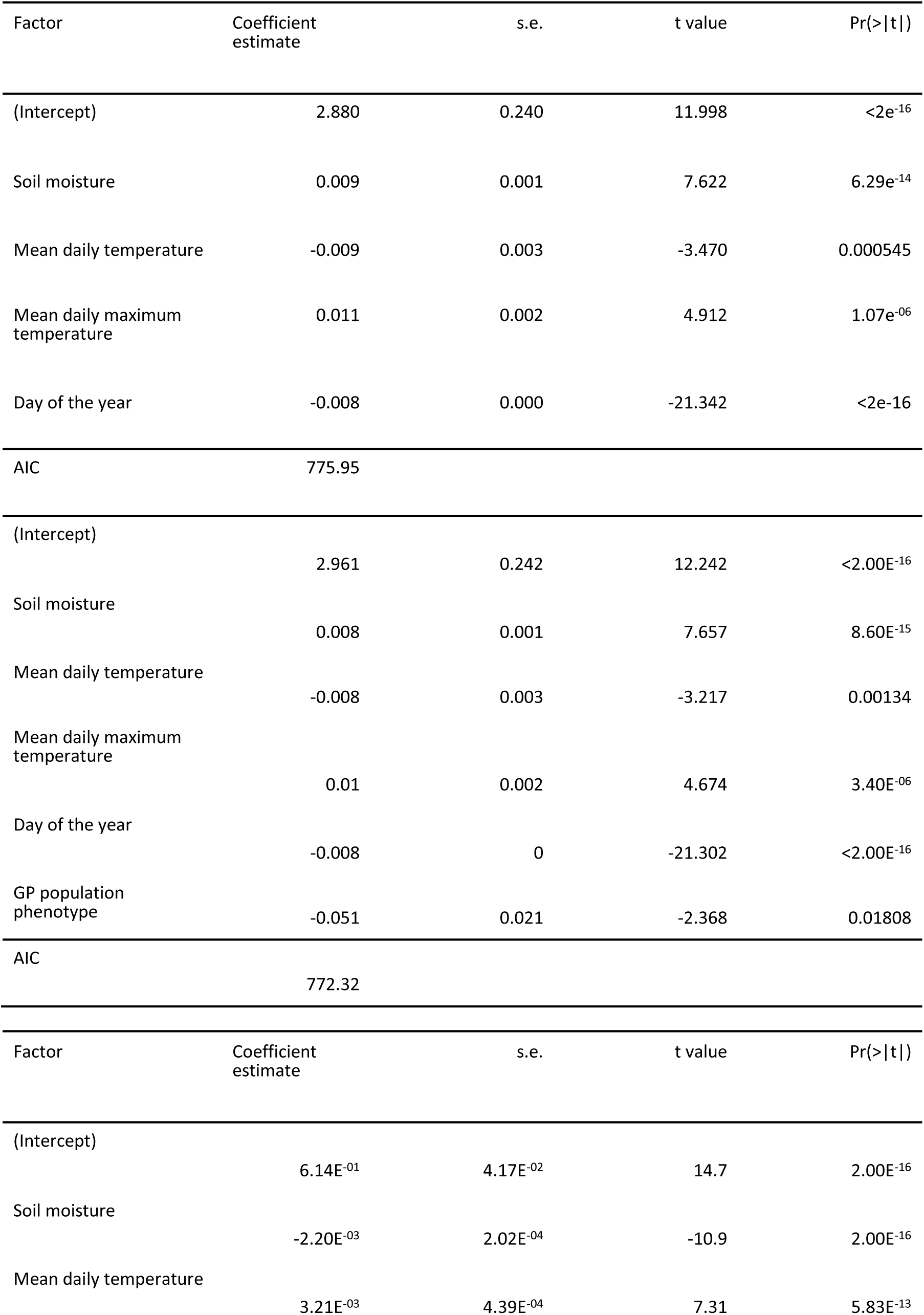
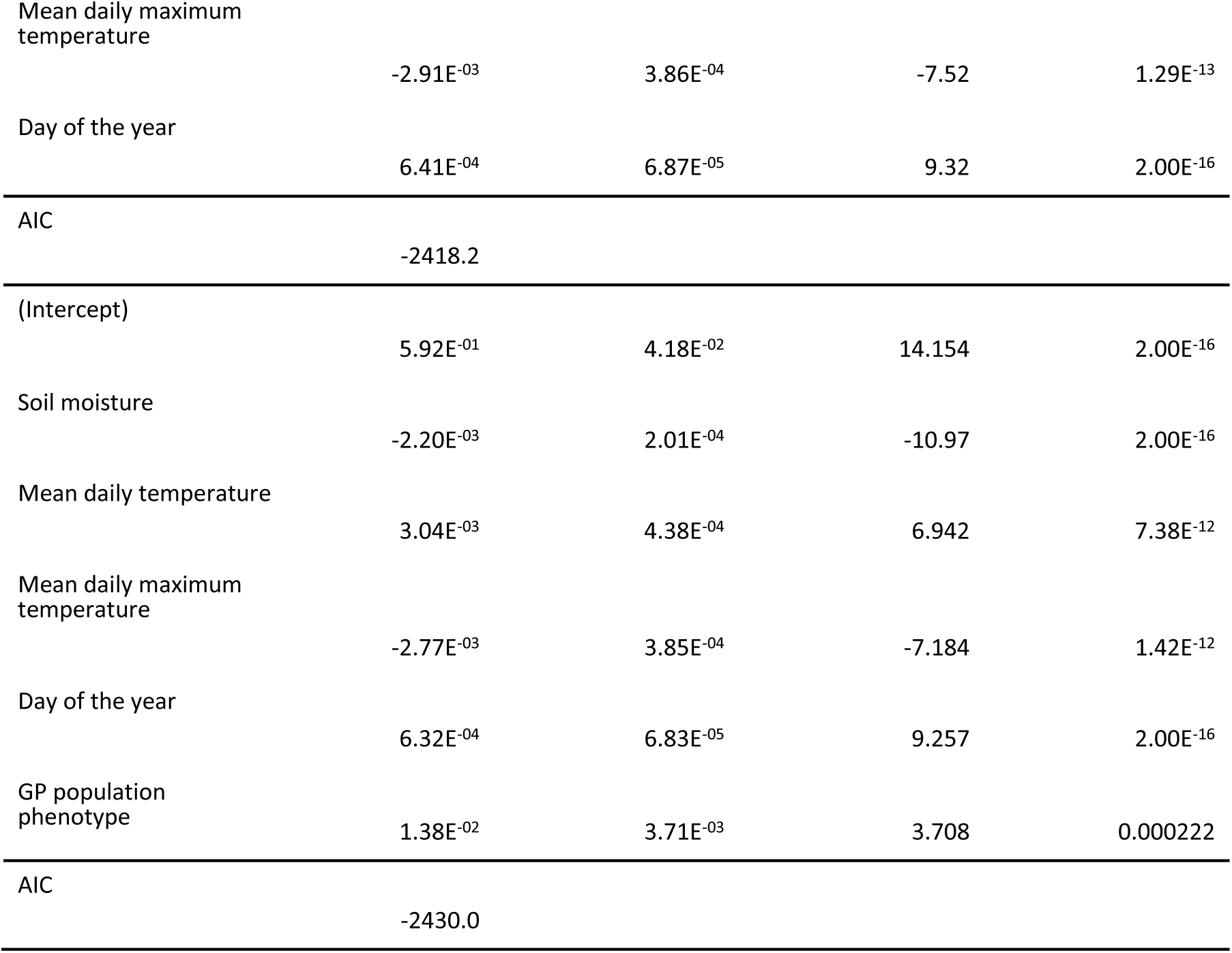
Model selection results. The upper table shows the results with Leaf Area Index (LAI) as response variable, below are the results for Moisture Stress Index (MSI).

### Predicted drought phenotype response to future climate scenarios

With no adaptation (PDPS unchanged) the long-term damage mean MSI threshold values of 0.625 will be regularly surpassed in the RCP 4.5 scenario for the decades 2041-2050 and 2091-2100 (Figure 4a). Under adaptation scenarios corresponding to i) an observed rate of phenotypic change (PDPS one unit lower) and ii) evolution to the level of the currently best adapted site (PDPS 0.599 at all sites), the expected climate conditions will not exceed the population’s drought tolerance (Figure 6). No adaptation scenario tested will likely allow persistence under the RCP 8.5 scenario (Supp. Fig. 3). The critical moisture stress threshold will be exceeded in any case in every year at all sites even in the decade 2041-2050.

**Figure 6.**
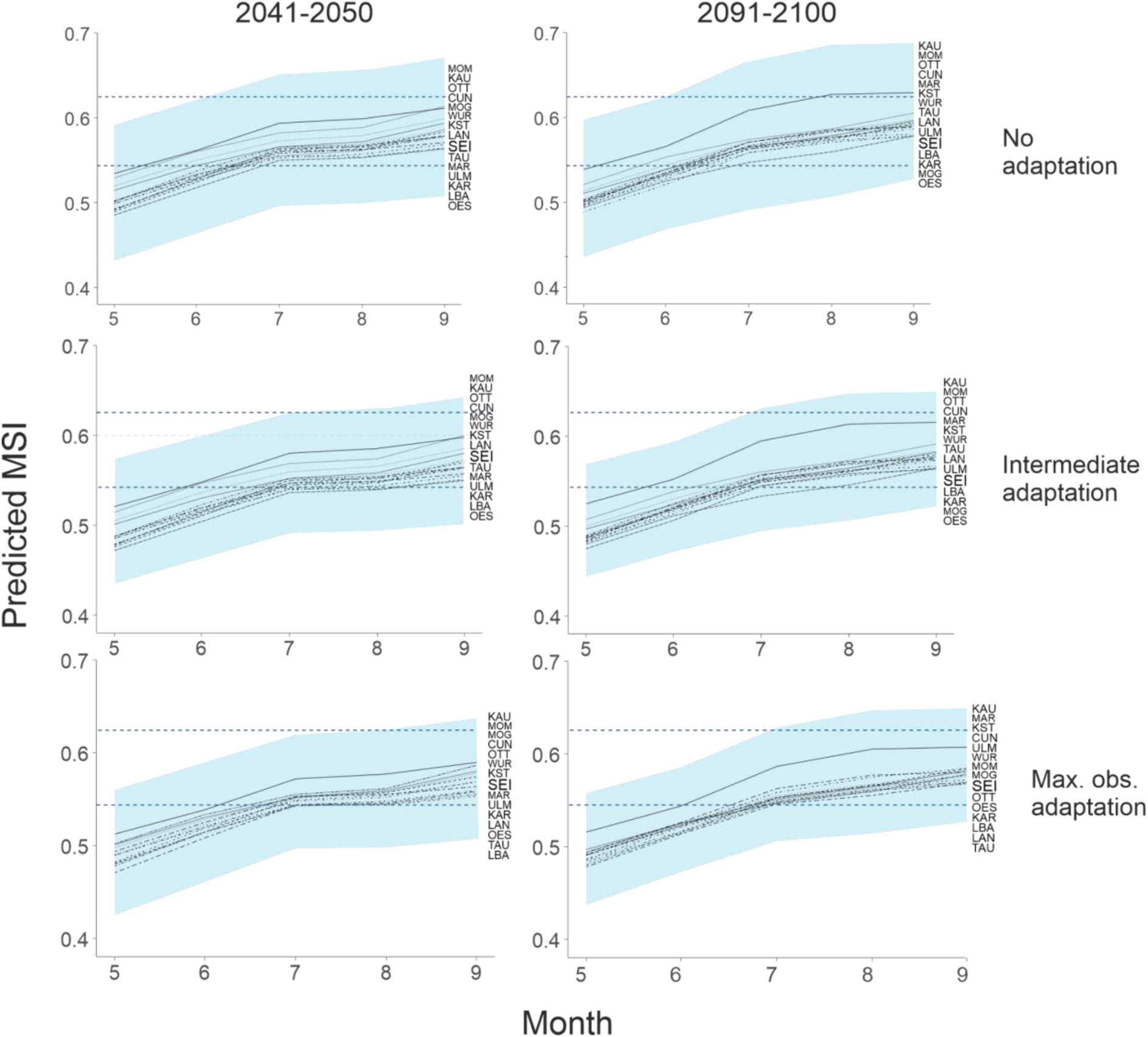
Predicted decadal mean MSI during the vegetation period (May-Sep) for future (2041-2050 and 2091-2100) climate scenario RCP 4.5 for all sites. The shaded area covers two standard deviations of the predicted weather variance and thus the expected inter-annual variation for the MSI range. The lower dashed blue line shows the stand mean MSI above which drought damage in leaf-cover can be observed. The upper dashed blue line indicates the lower mean MSI threshold for serious, long lasting damage of beech trees.

## Discussion

In this study, we integrated meteorological data, remote sensing and population genomics to predict the phenotypic response of European beech (*F. sylvatica*) forest tree stands to current and future climate dynamics. Our results show that their persistence under moderate climate change scenarios is possible if beech forests evolve based on existing genetic variation.

A validated genomic prediction model for the phenotypic drought response of beech individuals existed from a previous study, allowing the calculation of a continuous expected drought response score from the underlying multilocus genotype at associated loci ^14^. The current, population-based context required its extension to entire populations. As in the calculation of expected heterozygosity ^28^, estimated allele frequencies gained from the PoolSeq approach at predictive loci permitted calculation of a distribution of expected multilocus genotypes. The locus weighing vector derived from the individual genomic prediction model was then used to translate this distribution of multilocus genotypes into a distribution of predicted phenotypes. As with expected heterozygosity, this is technically the expected phenotype distribution in the zygotes of the next generation, but should also reflect the characteristics of the current population.

We compared the expected phenotype distribution derived from the allele frequencies over all stands with the range of actually observed values of trees in nature ^14^ to infer the processes underlying phenotypic evolution. The expected phenotype score distribution based on allele frequencies was wider than the observed range in the field. In particular individuals with increased drought susceptibility were missing in field individuals, suggesting stabilising selection on this trait ^29^. Since the alleles associated with the drought susceptible phenotype were often potential loss-of-function mutations ^14^, individuals with too many of them and thus high score values likely died before reaching maturity.

The predicted genetic component of phenotypic drought response varied substantially among stands. The observed PDPS ranged over almost two units; the largest difference was observed in the two most closely neighbouring stands, MOG and MOM. To illustrate the meaning of this difference, the expected proportion of trees not susceptible to drought ranged from 25 – 45%. As these simulated phenotype distributions are also an estimate of the expected progeny, the results showed that even at the currently worst genetically prepared site, a significant proportion of drought-resistant phenotypes should occur in the next generations. The partially large difference in PDPS among age classes (e.g. 1 unit or 0.41 Haldanes in the SH-class at LAN) suggested that a substantial shift in trait mean and thus better coping with future conditions is possible within a few decades, potentially as a result of ongoing natural selection. Overall, the results indicate that there is abundant standing genetic variation at the relevant loci at all sites. It thus seems likely that there is ample evolutionary potential for adaptation to more drought-resistant stands, especially considering the large number of offspring available for selection.

We could not detect any notable population structure among the *F. sylvatica* stands in the surveyed area. While the estimated mean F_ST_ of 0.039 was already quite low compared to other forest trees ^30–, 32^, this value is likely an overestimate as the comparison with the differentiation among growth classes within stands showed (Fig. 4). As the effective population size in *F. sylvatica* is large, drift should be negligible among contemporary generations ^33^. The observed level of genetic differentiation within stands is therefore mainly driven by sampling variance, i.e. the limited number of individuals used to obtain the estimate^34^. All stands sampled thus likely belong to the same, more or less panmictic, unstructured population. Absence of isolation-by-distance supported this view.

Reasons for this lacking population structure, also observed in other studies on beech ^35^, are likely wind pollination ^36^, partially over large distances ^37^, but also management practices that included long-range seedling and seed exchange ^38, 39^.

The effective absence of population structure, high genetic diversity, wind pollination by many partners over large spatial ranges, and extremely high number of offspring assure that most genetic diversity in *F. sylvatica* should be practically present everywhere all of the time. Low LD and high recombination rate quickly reshuffle beneficial (and deleterious) mutations by recombination in the many offspring. Intense demographic reduction from seed to seed ^40, 41^ with equally intense competition for light and space ^42^ will likely allow only individuals best coping with local conditions to reach canopy height and therefore permit swift adaptation in many independent trait dimensions simultaneously and, at the same time, ensure efficient removal of deleterious variation. One should therefore expect effective adaptation to the local conditions ^43^ in beech, as is expected for other forest tree species ^44^.

Despite these excellent preconditions, no local adaptation to drought risk was detectable in *F. sylvatica*. Neither the allele frequencies at underlying loci nor PDPS correlated with long term climate conditions predicting drought risk (Supp. Fig. 2). This could have two, mutually non-exclusive reasons. First, it is only in recent years that summer drought conditions may have regularly exceeded the limits of the phenotypic reaction norms allowing *F. sylvatica* to cope with environmental dynamics without loss of fitness. Given the multi-decadal generation time in beech, the period during which the increasingly changing selective climate regime is in effect has been too short for the canopy forming trees to adapt accordingly. Most of these beeches were already mature trees during the climate reference period 1970-2000. The climate during their youth and establishment phase was in most cases cooler and moister than this reference period ^14^. Even though serious drought years occurred occasionally (e.g. 1976), the current accumulation of these events is without precedent in recent beech generations. Moreover, a longer, stable, much cooler and wetter climate period, the Little Ice Age, lasting several beech generations ^45^, has likely shaped recent beech evolution ^46^, having rendered the population probably evolutionary unprepared for the current conditions. Second, many of the current canopy forming beeches were actively planted and raised in the past, thereby at least partially shielding the seedlings from natural selection, thus preventing local adaptation ^47^.

Our results showed that both crucial assumptions for GEA - a linear relation between relevant genetic and environmental/phenotypical variation and local adaptation - were not met in the present case. On the contrary, the search for the correlation of allele frequencies with either environmental or phenotypic variation likely would have produced much more false positives than truly associated loci, thus confirming recent findings ^18^. Also the assumption of local adaptation may not be taken for granted without proper validation ^16^. The framework introduced here could provide a feasible alternative to the currently favoured approaches. Phenotypic trait models as the proposed one have the advantage that they can be validated with contemporarily observed data in space and/or time.

Remote sensing offers an opportunity for objective, comparable long-term surveys of large geographical areas that would not be realisable otherwise. The satellite observations of beech stands were particularly valuable because they are a long-term measurement of reactions to environmental change and thus represent phenotypic reaction norms as a basis for modelling environmentally driven responses ^25^. Remote sensing observations of the stands, with each pixel likely corresponding to the canopy of individual trees, revealed a high spatial variability of response to the same weather conditions. This pattern was corroborated by previous visual observations of drought damaged trees often neighbouring apparently vital ones in forests ^14^. For useful evaluation of predictions under future climate scenarios, it was therefore necessary to find empirical threshold values of stand-wide mean MSI above which severe damage for parts of the trees occurs. Examining the temporal trajectories of individual pixels with large variance in LAI at sites with known damages by the 2018/19 droughts (e.g. LAN) suggested the existence of such thresholds (Figure 5c).

LAI values below 1.4 were almost never observed in any broad-leaf pixel in the non-drought baseline years 2016 and 2017. The percentage of pixels in a stand with values below this threshold appeared therefore to be a good indicator for the extent of drought-induced wilting or leaf-loss. Similar to the exemplary single pixel response (Figure 4D), LAI did not react to changes in MSI below a certain, site-independent threshold (0.544). Only above this value, moisture stress increasingly induced measurable leaf-damage. This suggests that this threshold is the phenotypic reaction norm limit to water stress in the surveyed beech population. While the vitality loss caused by mean MSI above 0.544 was reversible, values above 0.625 caused legacy effects that lasted over at least two years. Whether these legacy effects are reversible or were caused by partial canopy dieback or the entire tree, could not be inferred. Long-term vitality loss might be related to hydraulic system damage. Even though early leaf senescence in *F. sylvatica* is thought to protect the hydraulic system, loss of hydraulic conductance has been observed in the wake of the 2018 drought ^48^.

Despite the partial drought-induced vitality loss in the years after 2018 ^48^, the overall vitality status of surveyed *F. sylvatica* stands was mixed. LAI dropped below the critical damage threshold of 1.4 in up to 60% of the area at some sites during some observations and in one particularly drought-affected stand (LAN), more than 10 % of the area was damaged for a longer period of time (Figure 4E+F). However, a rather surprising result was that two-thirds of the total stand areas were never seriously affected by drought stress throughout the observation period, i.e. never had MSI values above 0.544.

Our approach was empirically confirmed by the fact that addition of PDPS explained the phenotypic variance in drought stress, measured as either LAI or MSI, significantly better than environmental variation alone. The genetic component explained with 11.4% (LAI), respectively 14.8% (MSI) a substantial part of the phenotypic variance. Only a small amount of variation (5.7%) was explained by systematic differences among the sampling sites. Unmeasured environmental differences among sites like e.g. soil composition, geological underground or biotic interactions ^11^ were therefore of minor importance for the phenotypic drought stress response.

Mean moisture stress experienced by the stands was mainly driven by the meteorological conditions, but the genetic component of the phenotype also played a substantial role. Obviously, PDPS reflects the population mean of the trait accurately enough to allow for efficient modelling the trait response to experienced weather conditions. This extension of genomic prediction to the logistically and economically efficient PoolSeq approach could be a major step forward for its use in climate change genomics ^49^.

Since moisture stress is the cause for changes in LAI under drought conditions ^11^ and environmental and genomic variation explained mean MSI better than mean LAI, we used MSI for predictive modelling. We used our validated model to predict the phenotypic response to drought to future climate scenarios. The choice of climate models, however, was restricted to those for which soil moisture as predicted parameter was available (RCP 4.5 and 8.5). We could show that already moderate, probably realistic rates of evolutionary adaptation based on standing genetic variation might be enough for most *F. sylvatica* stands to cope with a moderate climate change scenario such as RCP 4.5.

However, even though drought response is certainly crucial, other fitness-relevant traits, like e.g. phenology are also affected by the ongoing climate change ^50^. For a comprehensive assessment of the future development of beech forests, it will therefore be necessary to investigate these properties in a similar framework. But if both the tolerance limits and the evolutionary potential of drought resistance are likely to be exceeded in the future, as e.g. under the RCP 8.5 scenario, local extinction of the species is highly probable. Even though this worst-case scenario is not very likely to describe the future climate trajectory accurately, the threat of losing one of Europe’s most iconic ecosystem keystone species if the greenhouse gas emissions continue at the current rate should be motivation enough to act swiftly.

Persistence of beech forest in the study area under RCP4.5 and disappearance under RCP8.5 is consistent with results obtained with other methods. Buras and Menzel ^51^ combined an analysis of climate analogues with national forest inventory data, also using CMIP5 climate scenario data, but a larger set of Global Climate Models (GCMs). They found a decrease in relative abundance probability of *F. sylvatica* under RCP4.5 towards the end of the century and the disappearance of *F. sylvatica* in most of our study area under RCP 8.5 (Fig. 2 ^51^). Baumbach et al. ^52^ showed that future predictions from species distribution models for *F. sylvatica* across Germany also vary substantially between climate models (for the same RCP ^53^)

This study shows that the resistance of beech stands to drought is to a substantial part dependent on its genetic composition. It also shows that there is ample standing genetic variation for this trait at all sites investigated and therefore in the largely unstructured population as a whole. This suggests that natural selection by natural regrowth may be sufficient to maintain *F. sylvatica* as forest species in Central Europe, at least under moderate climate change scenarios. Whether assisted migration of potentially better adapted southern beech populations will provide additional advantages remains to be seen^54^, but might be beneficiary for timber production^8^. Approaches to replace native tree species with supposedly better adapted introduced species ^55^, however, could be premature as they could become disastrous for established ecosystems ^56^.

Under moderate climate change scenarios natural selection might act fast enough, but appropriately targeted studies and simulation models will be needed to confirm a sufficient evolutionary speed. The other, probably even more important condition is that future forest management practices employ close-to-nature forestry with several regeneration cycles to ensure a wide genetic basis allowing natural selection to work ^39^. An option to enhance natural selection in beech would be what we would like to term “Evolutionary Management”. The guiding principle here is to remove drought susceptible individuals as soon as possible or during regular thinning from the population to avoid their reproduction. Given that drought susceptibility has a genetic basis, selective logging would increase the proportion of drought resistant genotypes in the current and thereby also in the coming generations. At the same time, the capability for adaptation of the local population to other selectively relevant environmental dimensions would be preserved. While genomic prediction on individual trees would be feasible in principle, it would be costly and logistically challenging.

However, we have shown here that it is possible to identify drought sensitive patches in the stands with remote sensing early on as they respond more strongly to drought. Once it becomes possible to automatically separate and reliably identify individual trees from remote sensing data, drought susceptible trees could be removed selectively, even before they lose their economic value.

Rapid and widespread abiotic and biotic anthropogenically-driven changes of the earth system are occurring and will continue ^57^. In response to these changes, tools that enable fast and targeted management are required to maintain ecosystems. By coupling phenotyping with genomic and environmental data in key-trait focused phenotype models, we create a powerful and cost-effective tool with multiple potential applications. This approach is likely not only applicable to most tree species but has the potential to be extended to other ecosystem-shaping species. We foresee that such models integrating genomic, environmental, and phenotypic data could be used to anticipate climate change impacts on forests, pinpoint resilient forest stands, or speed the identification of less resilient stands allowing early intervention or proactive management. Moreover, such an approach can be combined with advanced technologies such as artificial intelligence or machine learning to enhance predictive capabilities and develop near real-time adaptive management strategies ^58^ that could enhance ecosystem resilience under changing environmental conditions.

## Material & Methods

### Sampled Fagus sylvatica sites

The sites selected for this study were in many regards typical for German beech forests ^6^. They comprised a large ecological amplitude with regard to long term conditions. Within the range from humid-cold to warm-dry climatic conditions important demographic and genetic processes in beech forests are taking place ^59^. Therefore, the aim of selection and stratification was to cover the climatic spectrum of forested areas in central and southern Germany to a large extent. For the characterisation of the climatic niche, we used the climatic marginality towards the rear edge ^60^, i.e. the dry and warm border of species distributions ^61^. A simple temperature increase scenario (+2.5 °C) was applied to estimate the site conditions to be expected in the future. The predicted climate was then divided into an optimal, intermediate, and marginal zone. Within the warm-dry zone we selected one stand on average soil condition (fresh water regime according to ^62^) and one stand with unfavorable water regime (future rear-edge). Besides the climate the nutrient regime (acidic, intermediate and basic soil) of the stands was taken into account. For each of the resulting 12 combinations, we established at least one representative site in a mature stand (minimum mean age of 70 years). As beech forest types in Germany are most frequent on acidic or intermediate soils ^6, 63^, these two strata were occupied by two stands (optimal/acidic: CUN, LAN; optimal/intermediate: ULM, WUR, Table 1). The more intensively studied stands KST and LAN are located at the edges of the climatic range at the warm-dry (marginal) and humid-cold (optimal) zones, respectively. All study sites were established in stands where beech had a share of at least 75 % of the total stand basal area. Forest stands with a history of frequent or heavy thinning were avoided to minimize potential management effects. All selected stands were larger than 4.5 ha. Thanks to the stratification scheme the full spectrum of beech forests with respect to climate and soil nutrient regime could be covered. The site condition of the most frequent beech forest types ^63^ were represented by two stands each. According to the Ellenberg Quotient (EQ) 4 sites are situated in the range (EQ 10-20) with beech forests having only low proportions of other tree species, 11 in the range (EQ > 20) where beech is usually admixed with other species, mostly oak. However, by including warm-dry sites on basic soils we also consider the range of a rarer type of beech forests containing a large biodiversity ^6^. The chosen stands comprised often unknown management histories with a mix of autochthonous and afforested sites which is typical for beech forests in Germany ^6^ and may have consequences for their evolution and adaptive capacity ^47^. We sampled 48 trees in a distance of 30 m to avoid the inclusion of closely related individuals. For the sites KST and LAN, two additional pools of 48 individuals from growth classes were sampled. The first class were trees below 1 m height (JW), the second established trees of 20-40 cm stem diameter in breast-height and not reaching the canopy (SH).

### Construction of population pools, sequencing, population structure and genetic diversity

From each tree 1-2 buds or 3-4 leave discs of 0.5 mm diameter (approx. 50 mg of fresh plant material) were dried in Silicagel prior to homogenization. DNA was extracted using an inhouse protocol. We constructed DNA pools per population using the same DNA quantity per individual beech. DNA concentration was measured using a Quantus fluorometer (Promega). Library preparation and 150bp paired-end sequencing with 450bp insert was conducted at Novogene.

Reads were trimmed using Trimmomatic v.0.39 ^64^ and quality controlled with FastQC v.0.11.9 ^65^. We used BWA mem v.0.7.17 ^66^ to map the reads onto the newest version of the beech reference genome ^67, 68^ and Samtools v.1.10 ^69^ to convert, sort and pile up the bam files. Duplicates were marked and removed with Picard v.2.20.8 (https://github.com/broadinstitute/picard). PoPoolation2 v.2.201 ^26^ pipeline was used to remove indels, and calculated allele frequencies for every position and pairwise FSTs in non-overlapping 1kb windows.

### Environmental data

To assess climatic differences among sites, we extracted long-term climatic data from Worldclim 2.1 data for the period between 1970-2000 with a resolution of 30 sec ^27^ for each sampling site using DIVA-GIS 7.5 ^70^. All temperature and precipitation parameters, including the BioClim parameters bio5 and bio18 were used. We summarised the data for the core vegetation period of beech at the sampling sites between May and September in a Principal Component Analysis (PCA).

Meteorological data on the actual weather conditions at the sampling sites during the core vegetation period between May and September was obtained for the years 2016 to 2022 from the DWD (Deutscher Wetterdienst) data portal at https://cdc.dwd.de/portal/202209231028/searchview. We used monthly grid data in a resolution of 1 x 1 km of the following parameters: mean air temperature, means of daily air temperature maxima and minima, sum of potential and real evapotranspiration, number of hot days (maximum >= 30 °C), number of summer days (maximum >= 25 °C), drought index according to de Martonne and medium soil moisture under grass and sandy loam. We summarised the data in a PCA.

### Genomic prediction from PoolSeq data

A recent article^14^ showed that individual genotypes at a selection of unlinked loci related to drought resistance can be successfully used to predict the respective phenotype of the tree based on a continuous phenotypic score in response to drought stress. Here we have extended this genomic prediction approach to PoolSeq data. In PoolSeq data, the information on individual genotypes in a population is lost in favour of precise allele frequency estimates ^71^. To take advantage of the existing genomic prediction framework, we used these allele frequency estimates at loci with high predictive power to simulate a large number of possible genotypes under the assumption of random mating within the population. We used the allele frequency information of 46 loci with high predictive power in linear discriminant analysis and the resulting weighting vector for each locus (Suppl. Table 1). For each sampling site, we simulated 10,000 individual genotypes. We then calculated the distribution of expected phenotype scores for each sampled population. This score is positive for susceptible phenotypes and negative for resistant ones. As the resulting distributions were approximately normally distributed, we used the mean as genomic predictor score of the population phenotype distribution of drought resistance (**P**opulation **<u>D</u>**rought **P**henotype **S**core, PDPS hereafter). The calculations of PDPS were performed with a custom Python script.

### Inference of local adaptation

If the populations at the sampling site were adapted to local drought conditions, we expected PDPS to be related to the long-term drought risk during the vegetation period. We therefore correlated the PCA1 of the long-term summer climate conditions (see above) to the PDPS of each site. To see whether the management history of the stands could have influenced the results, we fitted linear models to i) autochthonous stands and ii) stands known to be planted and of unknown origin separately. We also compared predicted phenotype score distributions among age classes at the sites KST and LAN.

### Remote sensing data

We focused on two remote sensing measurements of canopy features. Leaf Area Index (LAI; leaf area per ground area) and the Moisture Stress Index (MSI), that *a priori* seemed to best reflect the phenotype scored for genomic association (apparent vitality, respective loss of canopy leaves ^14^). Generally, the remote sensing values of LAI and MSI do not correspond directly to the canopy’s chemical, physical and biological characteristics but rather to the electromagnetic signal these produce. LAI is dimensionless and measures the covering of the ground area with leaves. The overlap of leaves can lead to a coverage of up to 16 under dense forest canopies ^72^. In beech forests, it ranges usually between 6 and 8. Remote sensing systems are not capable of capturing the heterogeneity in leaf distribution across forest canopies caused by this overlap. In this regard, retrieval methods commonly assume a simplified homogeneous canopy structure that makes the sensor actually measure a so-called ‘effective LAI’. It is described as the value that would induce the same remote sensing signal as the real LAI under the assumption of a random leaf distribution and is therefore inherently smaller ^73^.

In this study, we derived canopy LAI products of 10 m spatial resolution from atmospherically corrected Level-2 images acquired by the Multispectral Instrument on-board the ESA’s Sentinel-2 twin satellites. These products were accessed in the Sentinel Hub Earth Observation Browser (EO- Browser; https://www.sentinel-hub.com/explore/eobrowser/) for cloudless scenes during the growing season (May-Sep) from 2016 to 2022 using evaluation scripts. It allows the user to utilize the Sentinel-2 Level 2 Prototype Processor (SL2P) that is implemented in the Sentinel Application Platform (SNAP) software. The SL2P is an algorithm that enables the retrieval of vegetation biophysical variables from Level 2 optical satellite imagery. It is composed of artificial neural networks (ANNs) that were trained with simulations from the Leaf Optical Properties Spectra (PROSPECT) and Scattering by Arbitrarily Inclined Leaves (4SAIL) models. The ANNs are applied to top of canopy reflectance pixel values from nine bands between wavelength 560nm and 2190nm. In addition to the spectral information, acquisition geometry parameters (cosine of sun zenith angle, view zenith angle, relative azimuth angle) are being fed to the model ^74^. For the year 2016, canopy LAI products were generated using the same algorithm in SNAP based on Level-2A Sentinel-2 images accessed via the Copernicus Data and Exploitation Platform (CODE-DE).

MSI is, contrary to the LAI, an at-sensor parameter that is calculated as the ratio in near-infrared absorption at wavelength 819nm and 1599nm, and thus technically independent from LAI estimates. This ratio is a good indicator of water stress in plants, because the absorption of leaves in the vegetation cover at 1599 nm is positively correlated with their water content, while absorption at 819 nm remains almost unaffected. While usually ranging between 0.4 and 2.0, higher values are associated with greater water stress ^75^. In alignment to the retrieval of LAI, MSI was computed in the EO-Browser from images unaffected by cloud cover within the period of 2017 and 2021 over each study site by using an evaluation script that contains the respective index formula. For the year 2016, the MSI products were generated in QuantumGIS (QGIS) based on Level-2 imagery accessed from the CODE-DE platform.

### Damage threshold inference

Drought-induced leaf loss and thus decreased vigour is preceded by water (shortage) stress ^11^. To find empirical stand-wise mean MSI thresholds above which detectable drought-induced vitality degradation by leaf loss in at least some of the trees occurred, we first determined i) the 1% quantile of the minimum annual canopy LAI in the non-drought years 2016 and 2017 for each pixel at all sites, i.e. a lower limit for the normal LAI over the vegetation period (LAI_tres_) and ii) the mean maximum annual MSI for a pixel after which the above LAI_tres_ threshold was undercut for at least two years; i.e. that was followed by substantial, lasting leaf-loss (MSI_tres_). We then determined for all observation days and all stands the percentage of pixels that had values below LAI_tres_ or above MSI_tres_, respectively. These values were then regressed against the corresponding mean MSI of the entire stand. Threshold values were estimated from these plots by segmented regression ^76^.

### Model selection

To infer whether genomic prediction contributes significantly to the explanation of variance in LAI and MSI, we used general linear modelling (GLM) in a model selection approach. In a first step, we inferred the GLM that best explained the response variables with environmental data only. For each RS observation, the meteorological data (all variables listed above) for the respective month and the respective site were used as explanatory variables. We fitted GLMs to all possible parameter combinations and compared the associated Akaike information criteria (AIC). We considered the model with the lowest AIC to be the best model. We then added the PDPS as a further explanatory variable and calculated AIC for resulting models as well. All calculations were performed in R v… ^77^.

### Prediction of phenotypic reactions in future climate scenarios

Future climate conditions for each study site were derived from climate projections of the Climate Model Intercomparison Project 5 (CMIP5) that were bias-corrected and provided by the ISIMIP Fast Track Initiative. The datasets consist of five Global Climate Models (GCMs; HadGEM2-ES, IPSL-CM5A- LR, MIROC-ESM-CHEM, GFDL-ESM2M, and NorESM1-M) from which we calculated the assembled mean. We focused on two representative concentration pathways (RCP 4.5 and RCP 8.5). For each pathway and study site, we used time-series of daily mean air temperature and daily maximum air temperature for the next century from the ISIMIP data set, subset and aggregated to decadal means.

As a proxy for water stress, we used future projections of site-specific soil moisture given by an open-access dataset of future soil moisture index ^78^ based on simulations from the mesoscale Hydrologic Model (mHM; ^79, 80^), which was also calculated using the ISIMIP Fast Track. Again, soil moisture index was provided as five realisations, which were then aggregated to the ensemble mean. Finally, the soil moisture index of RCP 4.5 and RCP 8.5 were subset and aggregated to decadal means as the future temperature data.

### Introducing evolution in phenotypic prediction

We assumed three evolutionary adaptation scenarios: 1) No adaptation at all. This would correspond to a management shielding the stands completely from selection, e.g. by planting and raising random seedlings. 2) Intermediate adaptation. The PDPS of each site was diminished by the largest observed difference between two growth classes at the same site. Assuming that this difference was caused by natural selection, it would correspond to a natural adaptation rate that can be achieved within a few decades. 3) Maximum observed adaptation. In this scenario, we assumed that all sites evolve to the lowest PDPS observed. Given the genetic variation present in the surveyed *F. sylvatica* stands, more extreme PDPS and thus better drought adapted populations are theoretically possible, but it is uncertain whether such values are associated with trade-offs.

We used the decadal means for the two climate scenarios and the two periods to predict the MSI for each month in the vegetation period for each site. To account for the interannual variance in predicted weather conditions, we calculated a corridor of two standard deviations for the respectively most extreme sites.

## Data Availability Statement

Historical long-term climatic data from WorldClim database version 2.1 can be downloaded from https://www.worldclim.org/data/worldclim21.html, meteorological data from DWD can be accessed from https://cdc.dwd.de/portal/202209231028/searchview. Sentinel-2 surface reflectance data can be downloaded via the Sentinel hub services platform (https://apps.sentinel-hub.com/eo-browser/) and CODE-DE (https://finder.code-de.org/). Genomic data was archived at ENA https://www.ebi.ac.uk/ena/browser/home under the project accession number PRJEB60881.

## Acknowledgments

This study was partially funded by the Federal Ministry of Food and Agriculture and the Federal Ministry for the Environment, Nature Conservation, Nuclear Safety and Consumer Protection based on a resolution of the German Bundestag (project number 28W-B-4-058-01) and the Bavarian Office for Forest Genetics. We thank Enno Uhl and Matthias Steckel for support in the site and tree selection as well as Yves-Daniel Hofmann, Jorge Cueva-Ortiz, Friederike Reuss and Rebekka Stüwe for data acquisition in the field under sometimes very difficult conditions. Another part of funding was contributed by the Hessisches Landesamt für Umwelt, Naturschutz und Geologie (HLNUG) in the framework of the FAST-Project.

## Supplementary information

### Genomic prediction

**Supplementary table 1.**
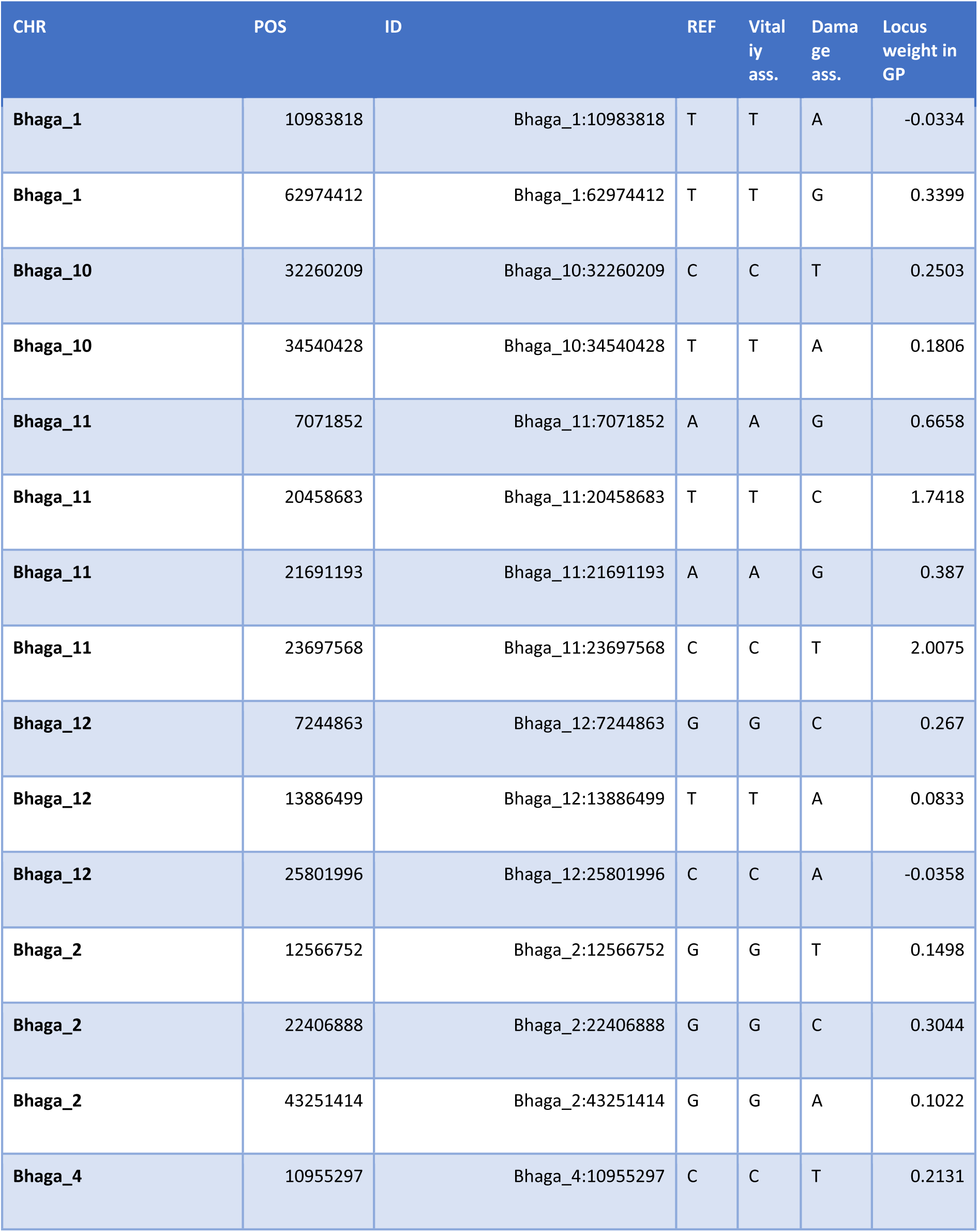
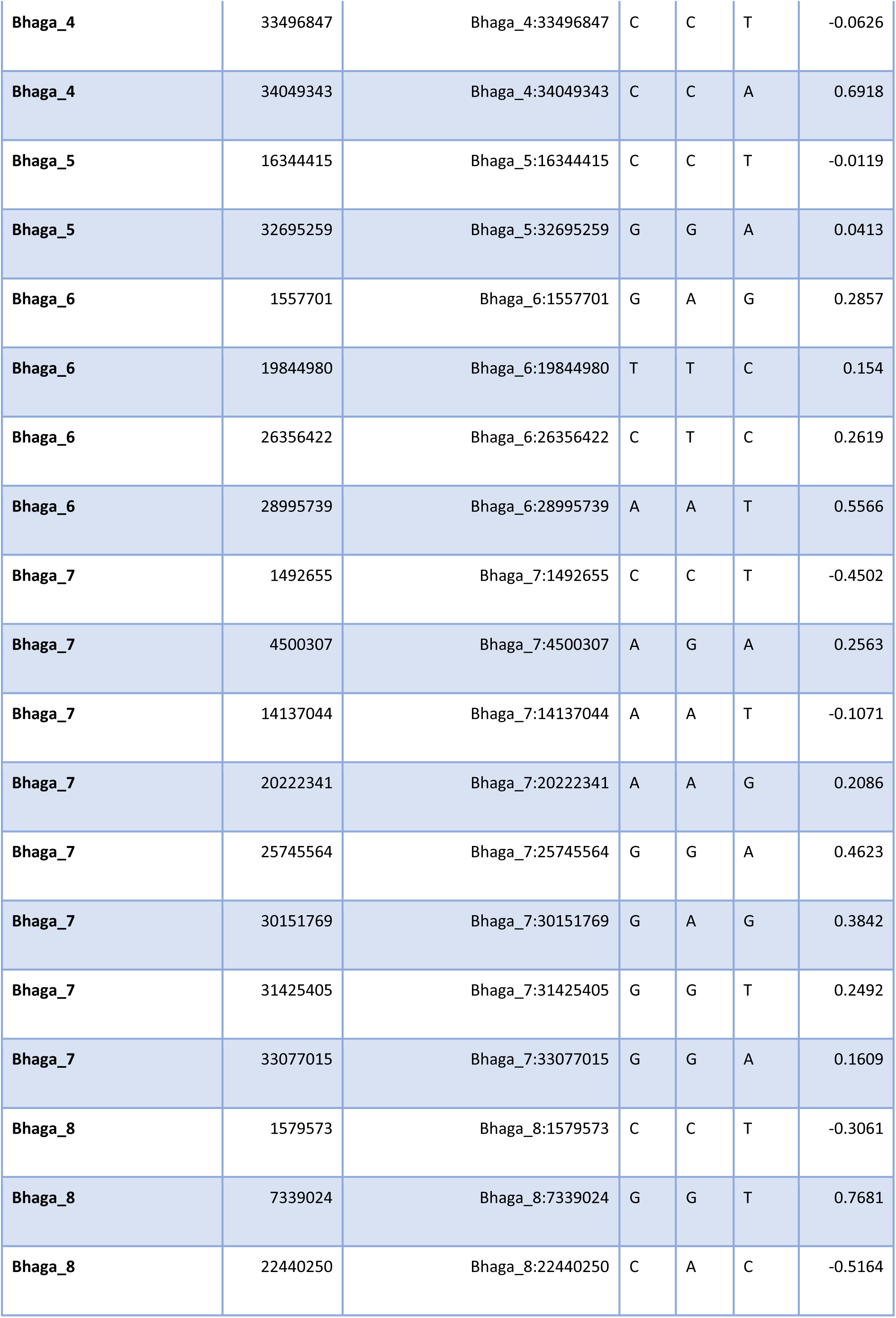
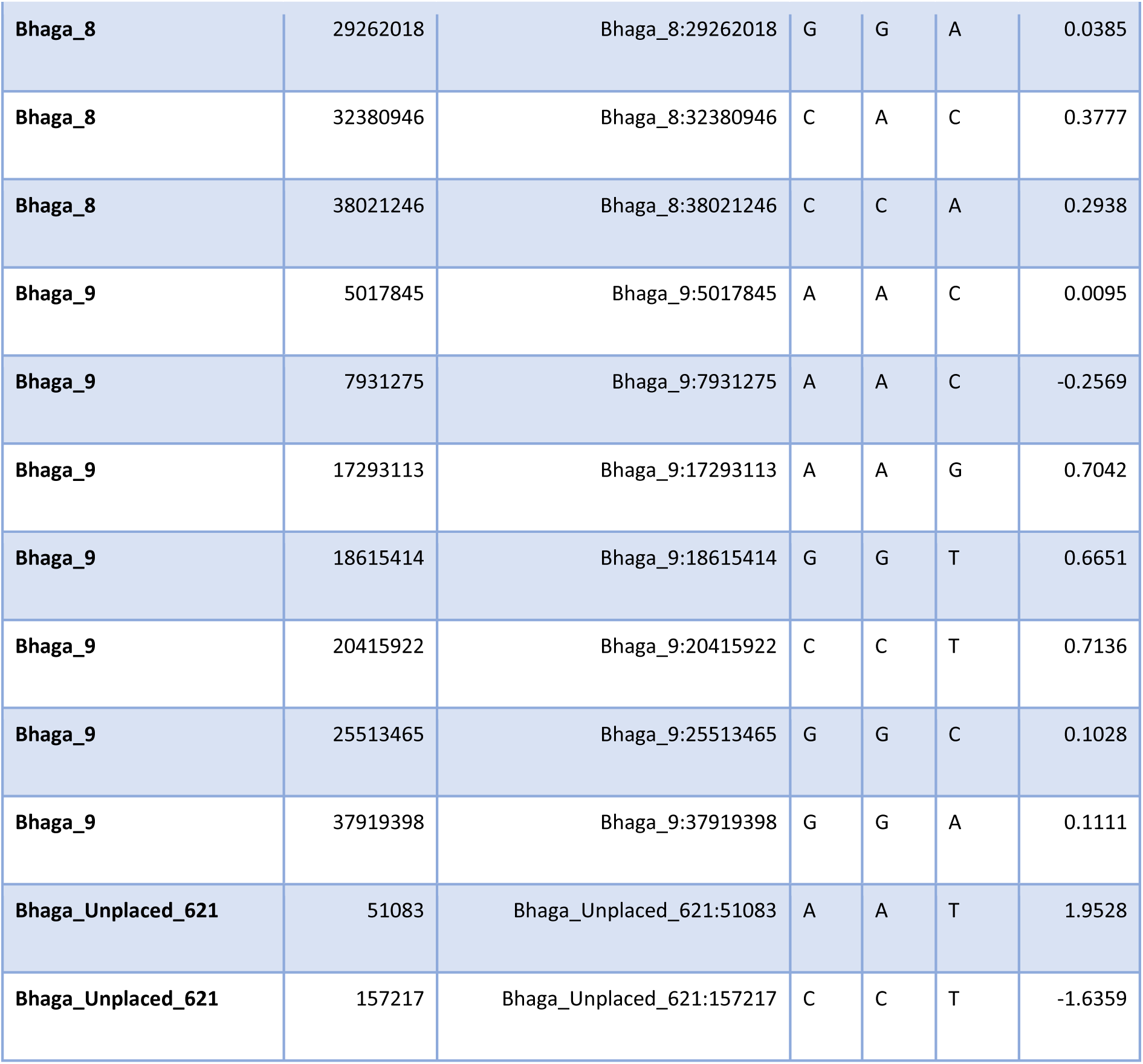
Loci used in genomic prediction, alleles associated with respective vitality conditions and genomic prediction (GP) weight.

| CHR | POS | ID | REF | Vital<br>ity<br>ass. | Dama<br>ge<br>ass. | Locus<br>weight in<br>GP |
| --- | --- | --- | --- | --- | --- | --- |
| Bhaga_1 | 10983818 | Bhaga_1:10983818 | T | T | A | -0.0334 |
| Bhaga_1 | 62974412 | Bhaga_1:62974412 | T | T | G | 0.3399 |
| Bhaga_10 | 32260209 | Bhaga_10:32260209 | C | C | T | 0.2503 |
| Bhaga_10 | 34540428 | Bhaga_10:34540428 | T | T | A | 0.1806 |
| Bhaga_11 | 7071852 | Bhaga_11:7071852 | A | A | G | 0.6658 |
| Bhaga_11 | 20458683 | Bhaga_11:20458683 | T | T | C | 1.7418 |
| Bhaga_11 | 21691193 | Bhaga_11:21691193 | A | A | G | 0.387 |
| Bhaga_11 | 23697568 | Bhaga_11:23697568 | C | C | T | 2.0075 |
| Bhaga_12 | 7244863 | Bhaga_12:7244863 | G | G | C | 0.267 |
| Bhaga_12 | 13886499 | Bhaga_12:13886499 | T | T | A | 0.0833 |
| Bhaga_12 | 25801996 | Bhaga_12:25801996 | C | C | A | -0.0358 |
| Bhaga_2 | 12566752 | Bhaga_2:12566752 | G | G | T | 0.1498 |
| Bhaga_2 | 22406888 | Bhaga_2:22406888 | G | G | C | 0.3044 |
| Bhaga_2 | 43251414 | Bhaga_2:43251414 | G | G | A | 0.1022 |
| Bhaga_4 | 10955297 | Bhaga_4:10955297 | C | C | T | 0.2131 |
| Bhaga_4 | 33496847 | Bhaga_4:33496847 | C | C | T | -0.0626 |
| Bhaga_4 | 34049343 | Bhaga_4:34049343 | C | C | A | 0.6918 |
| Bhaga_5 | 16344415 | Bhaga_5:16344415 | C | C | T | -0.0119 |
| Bhaga_5 | 32695259 | Bhaga_5:32695259 | G | G | A | 0.0413 |
| Bhaga_6 | 1557701 | Bhaga_6:1557701 | G | A | G | 0.2857 |
| Bhaga_6 | 19844980 | Bhaga_6:19844980 | T | T | C | 0.154 |
| Bhaga_6 | 26356422 | Bhaga_6:26356422 | C | T | C | 0.2619 |
| Bhaga_6 | 28995739 | Bhaga_6:28995739 | A | A | T | 0.5566 |
| Bhaga_7 | 1492655 | Bhaga_7:1492655 | C | C | T | -0.4502 |
| Bhaga_7 | 4500307 | Bhaga_7:4500307 | A | G | A | 0.2563 |
| Bhaga_7 | 14137044 | Bhaga_7:14137044 | A | A | T | -0.1071 |
| Bhaga_7 | 20222341 | Bhaga_7:20222341 | A | A | G | 0.2086 |
| Bhaga_7 | 25745564 | Bhaga_7:25745564 | G | G | A | 0.4623 |
| Bhaga_7 | 30151769 | Bhaga_7:30151769 | G | A | G | 0.3842 |
| Bhaga_7 | 31425405 | Bhaga_7:31425405 | G | G | T | 0.2492 |
| Bhaga_7 | 33077015 | Bhaga_7:33077015 | G | G | A | 0.1609 |
| Bhaga_8 | 1579573 | Bhaga_8:1579573 | C | C | T | -0.3061 |
| Bhaga_8 | 7339024 | Bhaga_8:7339024 | G | G | T | 0.7681 |
| Bhaga_8 | 22440250 | Bhaga_8:22440250 | C | A | C | -0.5164 |
| Bhaga_8 | 29262018 | Bhaga_8:29262018 | G | G | A | 0.0385 |
| Bhaga_8 | 32380946 | Bhaga_8:32380946 | C | A | C | 0.3777 |
| Bhaga_8 | 38021246 | Bhaga_8:38021246 | C | C | A | 0.2938 |
| Bhaga_9 | 5017845 | Bhaga_9:5017845 | A | A | C | 0.0095 |
| Bhaga_9 | 7931275 | Bhaga_9:7931275 | A | A | C | -0.2569 |
| Bhaga_9 | 17293113 | Bhaga_9:17293113 | A | A | G | 0.7042 |
| Bhaga_9 | 18615414 | Bhaga_9:18615414 | G | G | T | 0.6651 |
| Bhaga_9 | 20415922 | Bhaga_9:20415922 | C | C | T | 0.7136 |
| Bhaga_9 | 25513465 | Bhaga_9:25513465 | G | G | C | 0.1028 |
| Bhaga_9 | 37919398 | Bhaga_9:37919398 | G | G | A | 0.1111 |
| Bhaga_Unplaced_621 | 51083 | Bhaga_Unplaced_621:51083 | A | A | T | 1.9528 |
| Bhaga_Unplaced_621 | 157217 | Bhaga_Unplaced_621:157217 | C | C | T | -1.6359 |

**Supp. Fig. 1.**
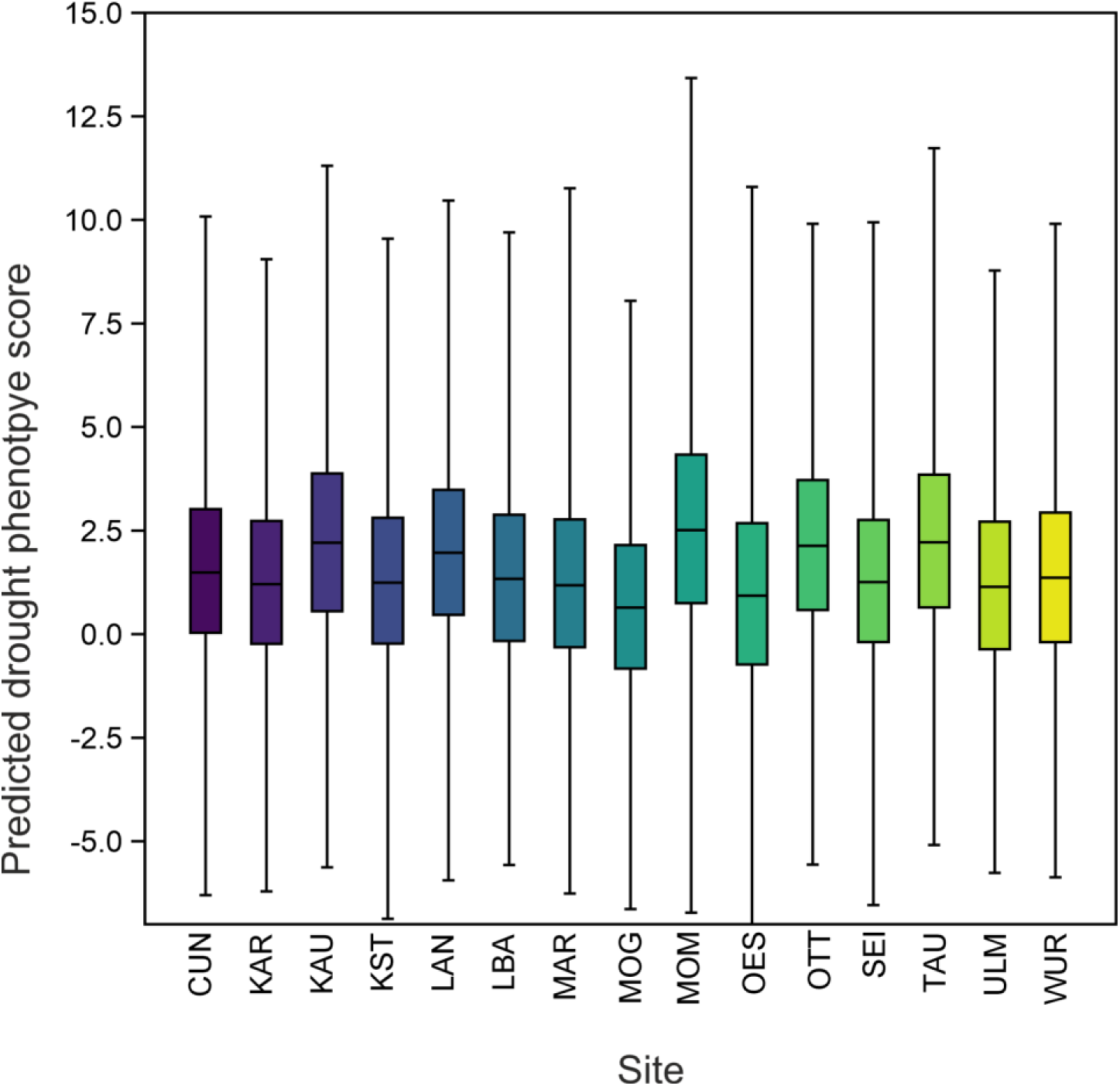
Box plot of individual drought phenotype score distributions estimated from the allele frequencies at 46 predictive loci for each site. The mean of this distribution corresponds to the population drought phenotype score PDPS.

**Supp. Fig.2.**
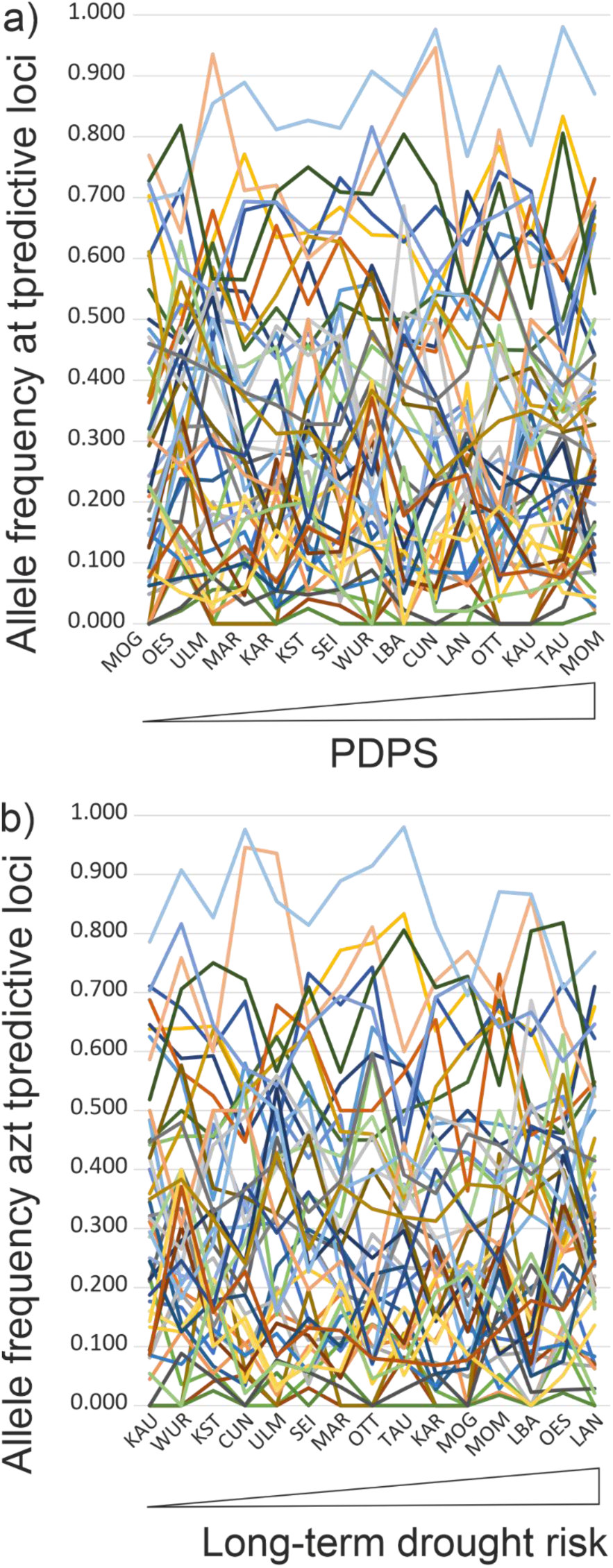
Allele frequencies at the 46 predictive loci for the fifteen sites. A) Sites ordered according to increasing predicted population drought phenotype score. B) Sites ordered according to their position on PCA1 on long-term summer climate conditions as drought risk predictor, drought risk increasing from left to right.

**Supp. Fig. 3.**
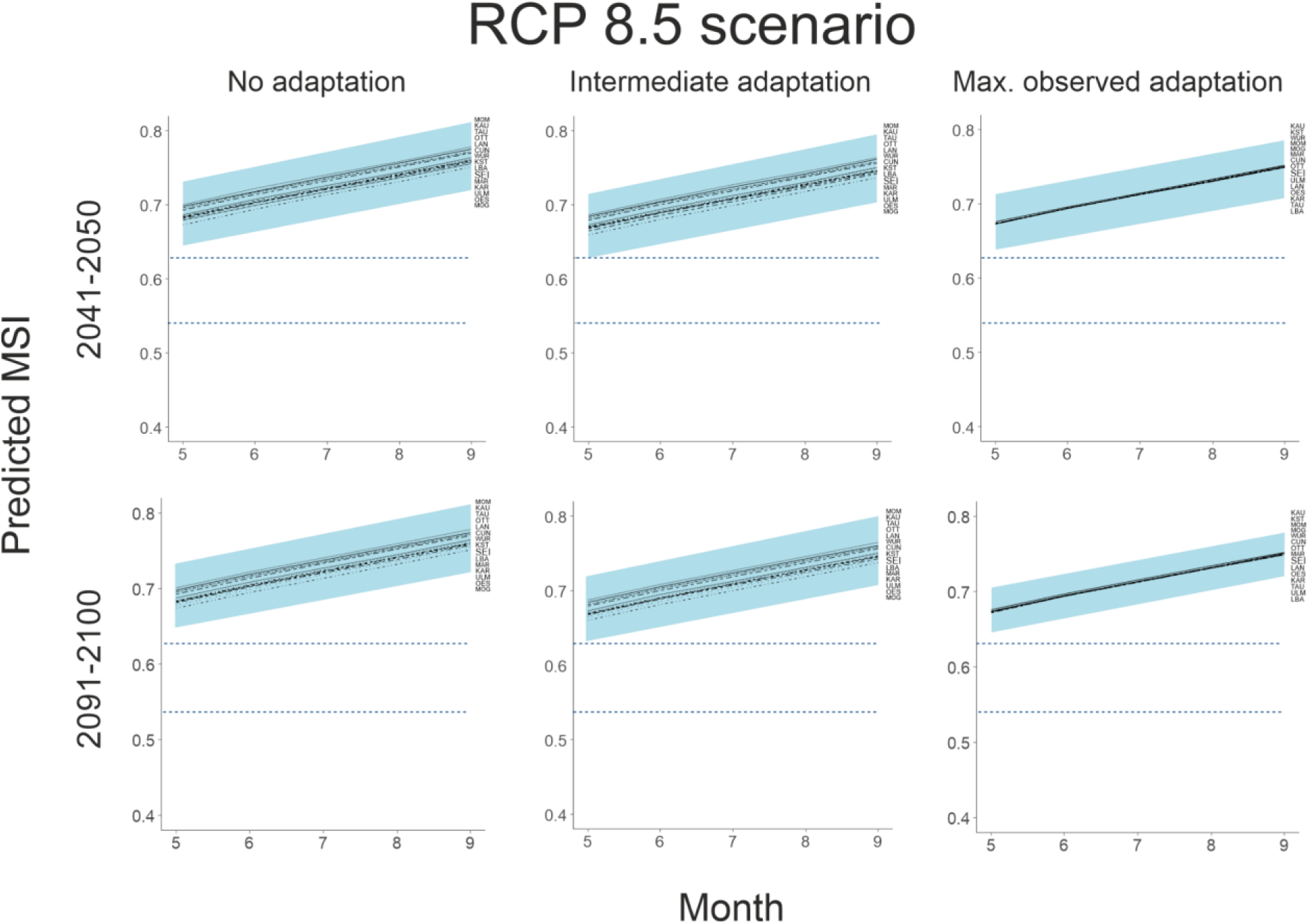
Predicted decadal mean MSI during the vegetation period (May-Sep) for future (2041- 2050 and 2091-2100) climate scenario RCP 8.5 for all sites. The shaded area covers two standard deviations of the predicted weather variance and thus the expected inter-annual variation for the MSI range. The lower dashed blue line shows the stand mean MSI above which drought damage in leaf-cover can be observed. The upper dashed blue line indicates the lower mean MSI threshold for serious, long lasting damage of beech trees.

